# An Evaluation of the Efficacy of Preclinical Models of Lung Cancer Drugs

**DOI:** 10.1101/2020.01.29.924779

**Authors:** Elizabeth Pan, David Bogumil, Victoria Cortessis, Sherrie Yu, Jorge Nieva

## Abstract

**Background:** Preclinical cell models are the mainstay in the early stages of drug development. We sought to explore the preclinical data that differentiated successful from failed therapeutic agents in lung cancer.

**Methods:** 134 failed lung cancer drugs and 27 successful lung cancer drugs were identified. Preclinical data were evaluated. The outcome variable for cell model experiments was the half maximal inhibitory concentration (IC50), and for murine model experiments was tumor growth inhibition (TGI). A logistic regression was performed on quartiles (Q) of IC50s and TGIs.

**Results:** We compared odds of approval among drugs defined by IC50 and TGI quartile. Compared to drugs with preclinical cell experiments in highest IC50 quartile (Q4, IC50 345.01-100,000nM), those in Q3 differed little, but those in the lower two quartiles had better odds of being approved. However, there was no significant monotonic trend identified (P-trend 0.4). For preclinical murine models, TGI values ranged from −0.3119 to 1.0000, with a tendency for approved drugs to demonstrate poorer inhibition than failed drugs. Analyses comparing success of drugs according to TGI quartile produced interval estimates too wide to be statistically meaningful, although all point estimates accord with drugs in Q2-Q4 (TGI 0.5576-0.7600, 0.7601-0.9364, 0.9365-1.0000) having lower odds of success than those in Q1 (−0.3119-0.5575).

**Conclusion:** There does not appear to be a significant linear trend between preclinical success and drug approval, and therefore published preclinical data does not predict success of therapeutics in lung cancer. Newer models with predictive power would be beneficial to drug development efforts.

## Background

Preclinical data guide the identification of oncology agents that have clinical promise^1^. However, the vast majority of agents with favorable preclinical data subsequently fail in human clinical trials. The cost to develop a new cancer drug ranges from $0.5 billion to $2 billion, and about 12 years typically elapse between selection of a candidate compound for human investigation to approval for clinical use. Drugs that enter human research use are met with a low ultimate FDA approval rate of 5-7%^2^, and there is a paucity of studies on whether satisfaction of preclinical criteria predicts eventual regulatory clearance. In regards to lung cancer drug development, specifically, a scholarly review published in 2014 (3) indicated that since 1998, only 10 drugs were approved for lung cancer treatment, while 167 other therapies failed in clinical trials^3^.

While mouse and cell models have elucidated pathophysiologic mechanisms of lung cancer, providing a biological framework for identification of therapeutic targets, new understanding that emerges from these efforts rarely translates into human therapeutics. Advantages of preclinical models include the far greater simplicity of both cell culture assays and animal model testing. By comparison, trials in humans are complicated by variability in patient factors such as genetic abnormalities, tumor microenvironment, metastatic potential in vivo, drug metabolism, and host immune responses. In addition, dosing schedules, drug delivery methods, and interactions between combination therapies vary significantly in humans compared to cell lines and murine models. These factors may account, at least in part, for failure of cancer therapies to achieve efficacy in clinical phase II and III trials. A recent study on oncolytic viral therapy illustrates the difficulty of applying *in vitro* success to clinical efficacy in humans. NTX-010, a picornavirus with selective tropism for small cell lung cancer tumor cell lines, and excellent preclinical data, was evaluated in a phase II study performed on 90 patients randomized to placebo versus treatment, and showed no benefit in progression-free survival in patients with small cell lung cancer^4^.

Similarly, many cancers have been cured in murine models but not humans^5^, illustrating limitations of preclinical testing in mice. It may be tempting to attribute these failures to the complexity and diverse evolutionary etiology of human cancers. However, even advanced cell-line derived xenografts and genetically engineered mouse models that produce tumors with great similarity to human diseases are not accurately and reproducibly translated to human applications.

In the work described here, we conducted in-depth review of design and results of preclinical cell and murine model experiments used in the development of lung cancer drugs, quantitatively comparing 27 drugs that are now FDA approved for treatment of lung cancer with 167 drugs that failed to be approved for this purpose. The goal was to identify features of preclinical experiments or values of efficacy parameters that might predict a drug’s success in clinical testing. Whether tested cells were of lung cancer origin was of particular interest, but any feature or efficacy measure found to be predictive could be emphasized to improve future preclinical testing in cells or animals. We recognized that should no such feature be identified, the analysis would underscore a need for alternate approaches.

## Materials and Methods

### Inclusion and exclusion criteria

We studied only drugs that had exhibited statistically significant efficacy in preclinical testing and subsequently entered the human testing phase of the United States Food and Drug Administration (FDA) approval process as candidates for single agent lung cancer therapy. From this set, we excluded any drug for which we could not determine specific model used in preclinical studies.

### Search strategy

We identified drugs that failed human testing using a PhRMA review of lung cancer medications that were unsuccessful in clinical trials from 1996-2014^3^. We identified approved drugs using the National Cancer Institute’s 2017 summary of medications approved by the FDA for treatment of lung cancer. We identified a corresponding set of preclinical studies, conducted either in cell lines or murine models, by systematically searching Pubmed through May 2018 using as search terms drug names taken from the lists described above together with the keywords, “lung cancer”, “preclinical mouse models”, “preclinical cell”, and “IC50”.

### Independent variables

For cell line experiments, the independent variable was the half maximal inhibitory concentration (IC50) expressed in nanomoles/liter (nM). This measure of efficacy is defined as the amount of drug needed to inhibit by half a specified biological process, which in these studies was cell growth.

The independent variable for mouse model experiments was tumor growth inhibition (TGI) calculated as (tumor volume or weight of treated mice in mm^3^ – tumor volume or weight of control mice in mm^3^)/tumor volume or weight of control mice in mm^3^ at the end of the follow-up period. TGI is 0 when the final size of tumors does not differ between drug-treated and vehicle-treated groups, <0 when drug-treated tumors are smaller, and >0 when drug-treated tumors are larger. For studies that used this definition of TGI, we used the reported value; if an alternative definition was used, we calculated the TGI according to the above formula from reported tumor volume and weight. For this purpose, we used Engauge Digitizer Version 10.4 application to estimate tumor volume or weight in treated and control mice.

We identified whether each drug was categorized as a nucleic acid damaging agent, cell signal-interrupting agent, tumor microenvironment and VEGF agent, immunotherapeutic agent (including vaccine), or miscellaneous (other). For cell culture models, we noted whether cells had been derived from lung cancer or non-lung cancer cell type. For animal models, we noted mouse strain categorized as athymic nude and immunocompetent, athymic nude only, or immunocompetent only; and coded tumor origin as xenograft, spontaneous, orthotopic implantation, induced, or murine vector.

### Outcome variables

The outcome variable for each analysis was drug approval status, scored as approved or failed.

### Statistical Analysis

To compare distributions of independent variables between failed and approved drugs, we created box-plots stratified by approval status. When raw data were highly skewed, we log transformed IC50 values and created a second set of box-plots on this scale. To test for differences in central tendency, we used T-tests for normally distributed data and the Mann-Whitney procedure for skewed data, and reported p-value results of each.

We used logistic regression to estimate associations between drug approval status and quartile of IC50 (cell studies) or TGI (animal studies), and calculated trend P-values based on IC50 or TGI value of midpoint of each quartile. We estimated conventional standard errors of TGI. Since there were numerous cell studies of some drugs, we recognized that there could be dependence between measures and thus employed generalized estimating equations to estimate robust standard errors of IC50 to accommodate this apparent non-independence.

Finally, we created empirical receiver operator characteristic (ROC) curves displaying sensitivity and specificity of each value of the independent variable to predict a drug’s success. We created a single ROC curve for TGI values; for IC50 values, we created one curve for all measures, and separate curves for studies that employed cell lines derived from lung cancer or from other tissues.

All analyses were conducted using R 3.5.1.^6^

## Results

Our search identified reports on preclinical studies of 161 drugs that had been carried forward to human testing as part of the FDA approval process. Of these, 27 had been approved as monotherapy for lung cancer, but 134 had failed at some stage of human testing.

### Preclinical cell models

Our search identified reports on 378 cell culture experiments (Supplemental Table 1). IC50 values were not reported for 55 of these, precluding their use in the analyses. Table 1 summarizes the remaining 323 experiments according to type of drug and cell line used, and provides of IC50 values that define each quartile of this variable for failed and approved drugs. Cell lines derived from lung cancer were used in only 25% of experiments that tested approved drugs and 17.3% of studies of drugs that failed.

**Table 1:**
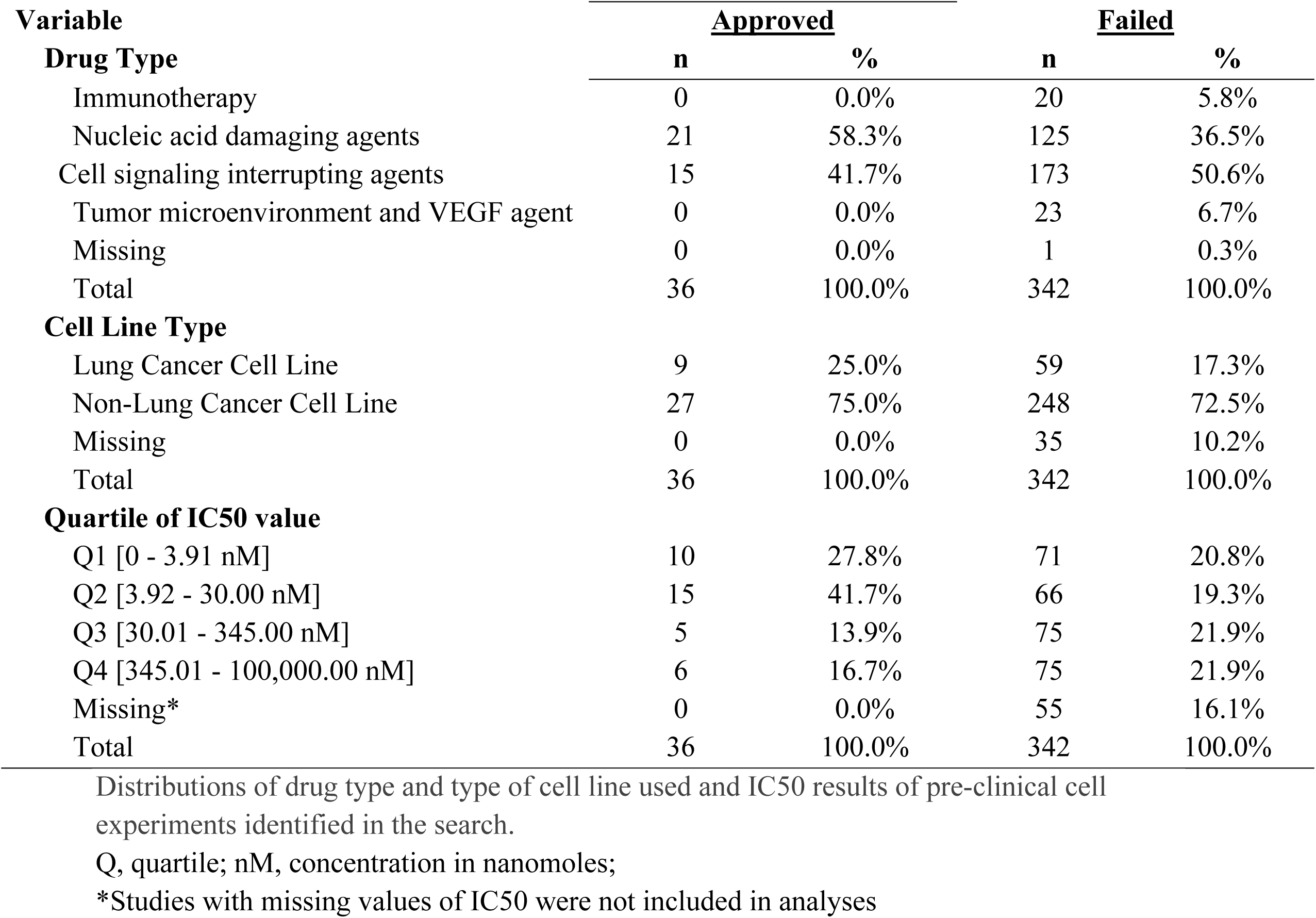
Descriptive Distributions on Initial Sample.

| Variable | <u>Approved</u> |  | <u>Failed</u> |  |
| --- | --- | --- | --- | --- |
|  | n | % | n | % |
| **Drug Type** |  |  |  |  |
| Immunotherapy | 0 | 0.0% | 20 | 5.8% |
| Nucleic acid damaging agents | 21 | 58.3% | 125 | 36.5% |
| Cell signaling interrupting agents | 15 | 41.7% | 173 | 50.6% |
| Tumor microenvironment and VEGF agent | 0 | 0.0% | 23 | 6.7% |
| Missing | 0 | 0.0% | 1 | 0.3% |
| Total | 36 | 100.0% | 342 | 100.0% |
| **Cell Line Type** |  |  |  |  |
| Lung Cancer Cell Line | 9 | 25.0% | 59 | 17.3% |
| Non-Lung Cancer Cell Line | 27 | 75.0% | 248 | 72.5% |
| Missing | 0 | 0.0% | 35 | 10.2% |
| Total | 36 | 100.0% | 342 | 100.0% |
| **Quartile of IC50 value** |  |  |  |  |
| Q1 [0 - 3.91 nM] | 10 | 27.8% | 71 | 20.8% |
| Q2 [3.92 - 30.00 nM] | 15 | 41.7% | 66 | 19.3% |
| Q3 [30.01 - 345.00 nM] | 5 | 13.9% | 75 | 21.9% |
| Q4 [345.01 - 100,000.00 nM] | 6 | 16.7% | 75 | 21.9% |
| Missing* | 0 | 0.0% | 55 | 16.1% |
| Total | 36 | 100.0% | 342 | 100.0% |

Distributions of drug type and type of cell line used and IC50 results of pre-clinical cell experiments identified in the search.
Q, quartile; nM, concentration in nanomoles;
\*Studies with missing values of IC50 were not included in analyses

Reported IC50 values range from 1nM to 100,000nM, with substantial overlap in distributions within the set of drugs that were approved and those that failed. Values for approved drugs were slightly lower than values for drugs that failed, but the difference did not achieve statistical significance (means of log(IC50), p=0.22; medians of IC50, p=0.09; Fig 1). Accordingly, estimated areas under the ROC (AUC) values were only slightly greater than 0.5, consistent with IC50 predicting success barely better than chance, whether the ROC represented data from all preclinical cell experiments (AUC=0.59, Fig 2A) or from subsets (Fig 2B) defined by whether the cell line originated from lung cancer (LCLine, AUC=0.56) or some other source (non-LCLine, AUC=0.60).

**Fig 1.**
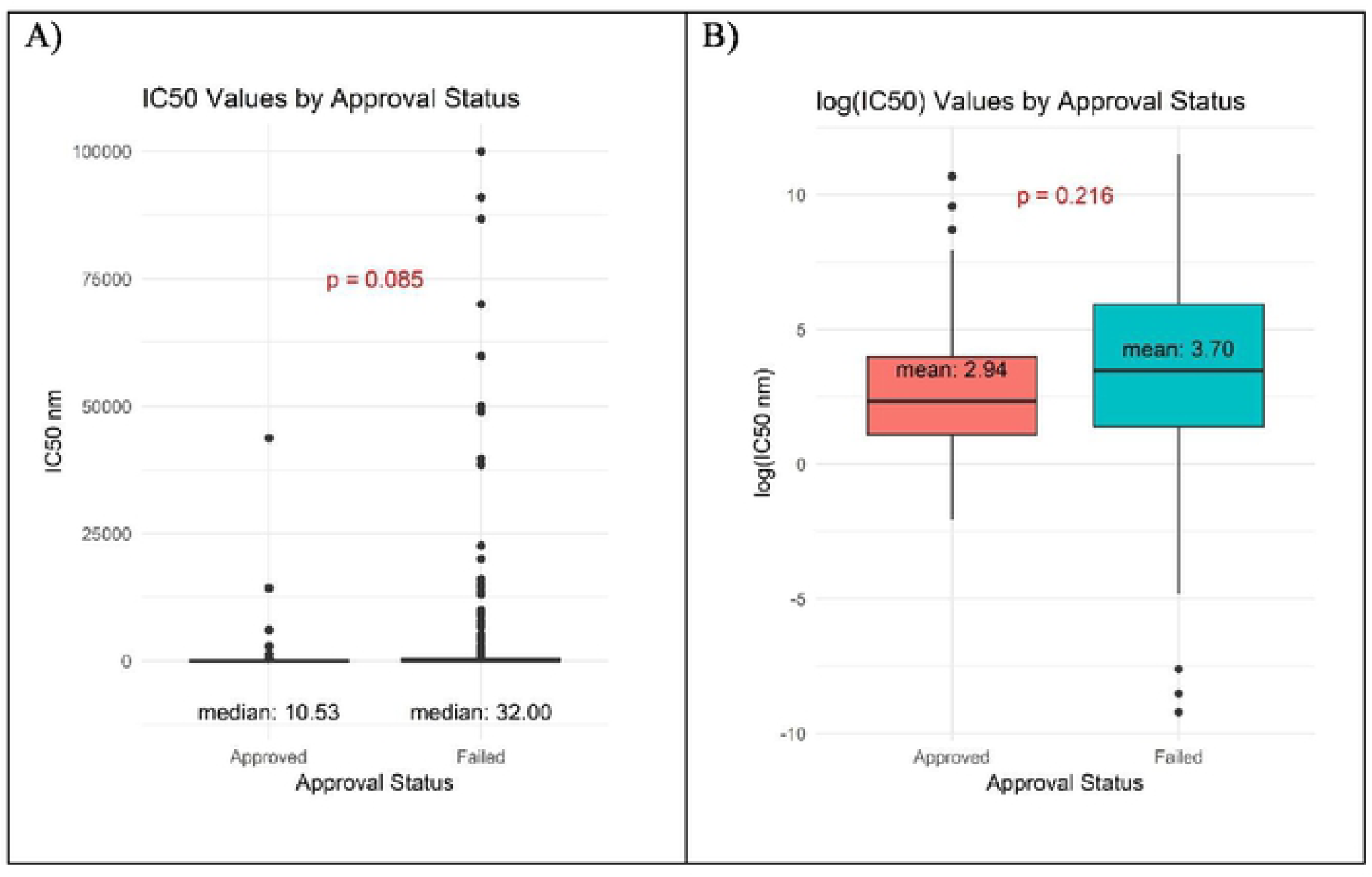
Distributions of IC50 values in preclinical cell line experiments among drugs that were subsequently approved or failed. A) IC50 values and B) log(IC50).

**Fig 2.**
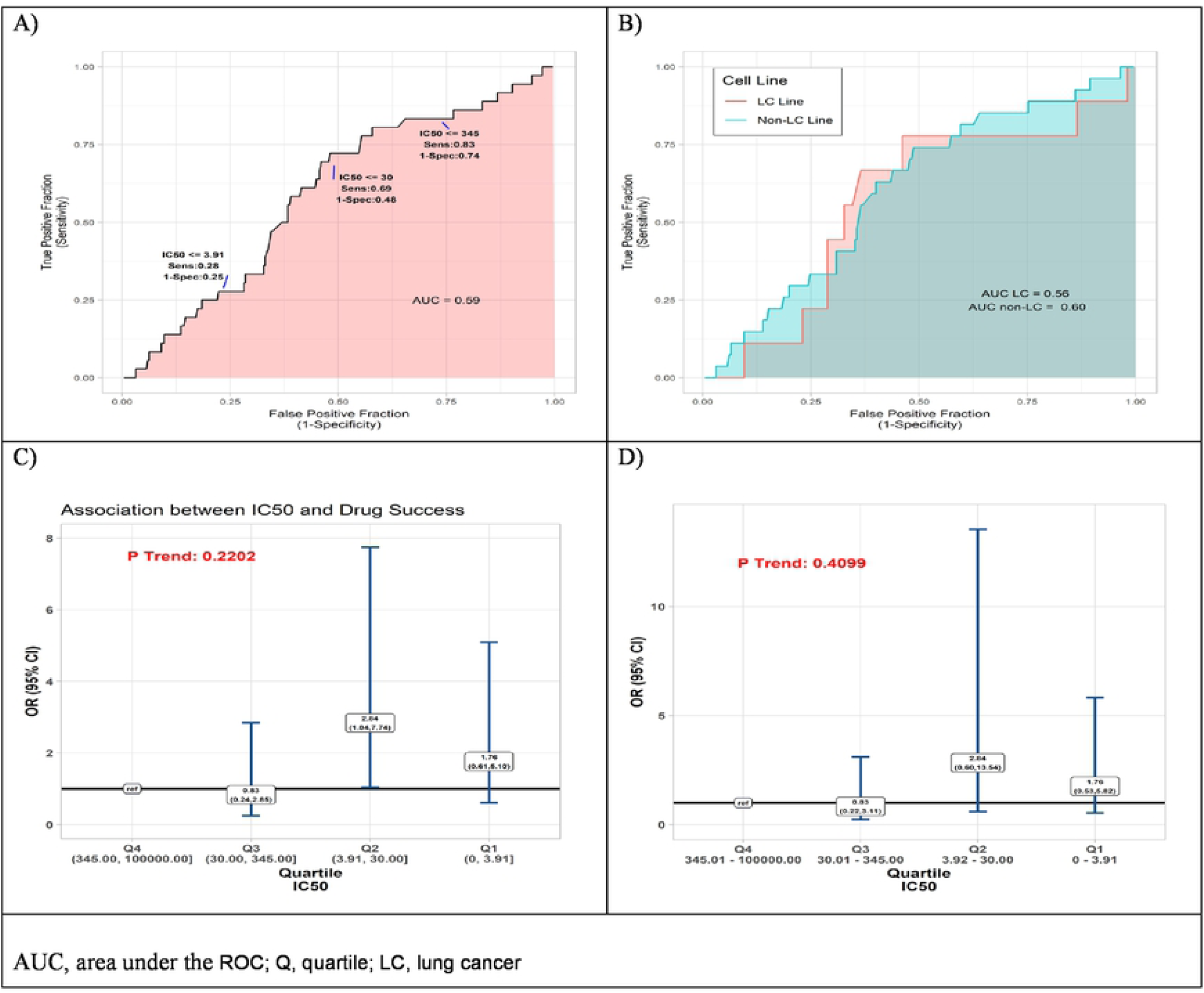
Results of quantitative analyses of preclinical cell line experiments. A) Receiver operator curves (ROC) displaying accuracy of IC50 value as predictor of drug approval for all cell line experiments combined, and B) within subsets defined by type of cell line, lung cancer cell lines (pink) non-lung cancer cell lines (aqua). C) Odds Ratio (OR) associations between drug success and quartile of IC50 result of pre-clinical cell model experiment, drugs with IC50 in lower quartiles (Q) Q1, Q2, Q3 compared to those with IC50 in highest quartile, Q4 (reference) by two analytic methods, conventional logistic regression, and D) General Estimating Equation (GEE), allowing for non-independence of multiple experiments using the same drug.

In a final set of analyses of these data, we compared odds of approval among ordinal categories of drugs defined by IC50 quartile. Compared to drugs in the highest quartile (Q4, IC50 345.01-100,000nM), those in the third quartile differed little, but those in the lower two quartiles had somewhat better odds of being approved. Most favorable results were for drugs in the second quartile (Q2, IC50 3.92-30nM) for which the estimate from conventional logistic regression was OR=2.84 (95%CI 1.04-7.74). However, results from the more conservative GEE analysis – which accounts for possible non-independence of results from multiple experiments using the same drug – do not achieve statistical significance (OR=2.84 [95%CI 0.60-13.54]). Neither analysis identified a statistically significant monotonic trend in effect size (Fig 2C-D).

### Preclinical murine models

The search identified 144 preclinical studies using murine models of lung cancer drugs that satisfied inclusion criteria, with all published reports providing sufficient experimental data to use in our analyses (Supplemental Table 2). The measure of efficacy used in these experiments was TGI. Table 2 summarizes the studies according to type of drug and mouse model, TGI measure employed, and quartile of TGI efficacy among results of all studies.

**Table 2.** Distributions of preclinical animal study data (n = 213).

|  | Approved |  | Failed |  |
| --- | --- | --- | --- | --- |
|  | n | % | n | % |
| **TGI\* measure types:** |  |  |  |  |
| Tumor Volume | 28 | 66.7% | 89 | 52.0% |
| Tumor Weight | 5 | 11.9% | 12 | 7.0% |
| Other | 3 | 7.1% | 11 | 6.4% |
| Missing | 6 | 14.3% | 59 | 34.5% |
| Total | 42 | 100.0% | 171 | 100.0% |
| **Quartile of TGI** |  |  |  |  |
| Q1 [-0.3119, 0.5575] | 10 | 23.8% | 26 | 15.2% |
| Q2 [0.5576, 0.7600] | 9 | 21.4% | 28 | 16.4% |
| Q3 [0.7601, 0.9364] | 9 | 21.4% | 26 | 15.2% |
| Q4 [0.9365, 1.0000] | 8 | 19.0% | 28 | 16.4% |
| Missing | 6 | 14.3% | 63 | 36.8% |
| Total | 42 | 100.0% | 171 | 100.0% |
| **Drug Type** |  |  |  |  |
| Immunotherapy (including vaccines) | 0 | 0.0% | 11 | 3.7% |
| Nucleic acid damaging agents | 18 | 42.9% | 46 | 28.7% |
| Cell signaling interrupting agents | 24 | 57.1% | 84 | 50.0% |
| Tumor microenvironment and VEGF agents | 0 | 0.0% | 29 | 17.6% |
| Missing/NA | 0 | 0.0% | 1 | 0.0% |
| Total | 42 | 100.0% | 171 | 100.0% |
| **Mouse Type** |  |  |  |  |
| Xenograft on nude mouse | 42 | 100.0% | 148 | 94.4% |
| Spontaneous tumor model | 0 | 0.0% | 4 | 2.8% |
| Orthotopic model (same origin site of tumor) | 0 | 0.0% | 5 | 2.8% |
| Induced tumor model (chemical, radiation, genetic, etc) | 0 | 0.0% | 1 | 0.0% |
| Missing/NA | 0 | 0.0% | 13 | 0.0% |
| Total | 42 | 100.0% | 171 | 100.0% |

\*TGI, tumor growth inhibition, a measure of efficacy estimated as described in Methods

TGI values ranged from −0.3119 to 1.0000, with a tendency for approved drugs to demonstrate slightly poorer inhibition than drugs that failed to be approved. The respective medians were 0.74 and 0.77, a small difference that does not achieve statistical significance (P=0.375, Fig 3A). Analyses comparing success of drugs according to quartile of TGI produced interval estimates too wide to be statistically meaningful, although all point estimates accord with drugs in each of the three highest quartiles (Q2-Q4, TGI 0.5576-0.7600, 0.7601-0.9364, 0.9365-1.0000) having lower odds of success than those in lowest quartile (Q1, -0.3119-0.5575) (Fig 3B). In accordance with these results, the AUC estimate was 0.45, (Fig 3C), corresponding to TGI value performing slightly worse than chance for predicting eventual success of a drug.

**Fig 3.**
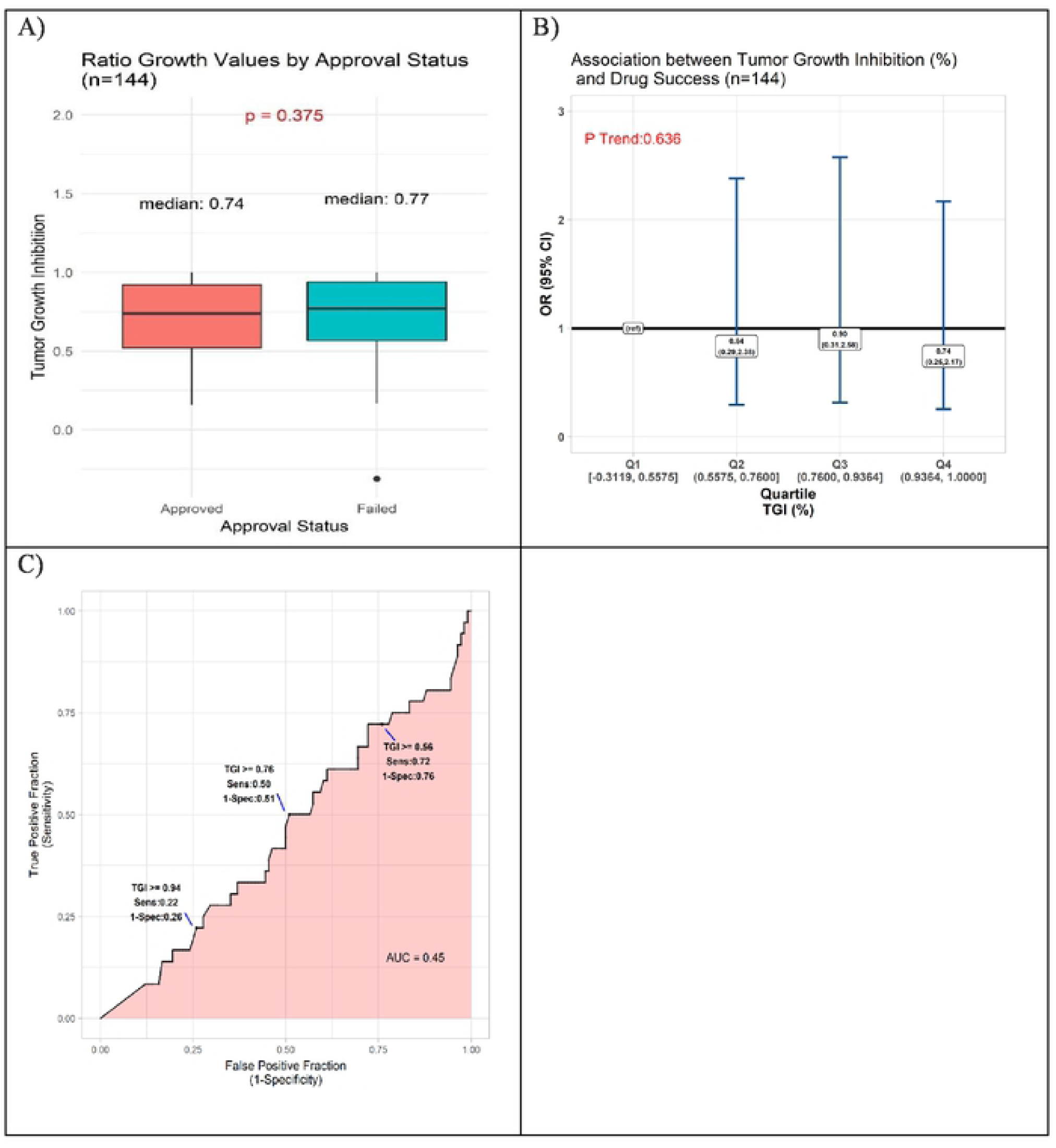
Results of quantitative analyses of preclinical studies in murine models. A) Box-plots displaying distributions of tumor growth inhibition (TGI) in preclinical murine models of subsequently approved and failed drugs. B) Odds Ratio (OR) estimates of association between drug success and TGI result of pre-clinical animal model experiments, drugs with TGI in each of quartiles (Q) Q1, Q2, Q3 compared to those with TGI in Q4 (reference). C) Receiver operator curve displaying accuracy of TGI value as predictor of drug approval.

## Discussion

We endeavored to quantitatively investigate predictive value of publicly available results from preclinical studies of lung cancer drugs, conducted over nearly two decades. This novel effort identified no value of efficacy parameters that predicted approval of lung cancer drugs.

The current FDA guidelines require animal testing prior to human exposure^7^, with the hope that preclinical results may be mimicked in human subjects. Unfortunately, most successful preclinical testing falls short of expectations, with only a third of preclinically approved drugs entering clinical trials^8^ at a failure rate of 85% (all phases included), and a 50% success rate in the fraction of therapeutic agents that make it past phase III^9^. Anti-cancer agents account for the largest proportion of these failures^10^. Flawed methodologies in clinical trial testing may be contributing to the disparity in preclinical and clinical success. Additionally, it is estimated that animal studies overestimate by 30 percent the likelihood of treatment efficacy due to unpublished negative results^11^. The poor positive predictive value of successful preclinical testing has been attributed largely to disparity between disease conditions in mice and humans. The nature of the animal model and laboratory conditions, which are currently not standardized, may also contribute to variations in animal responses to therapeutic agents^12^.

There have been several published examples of successful cancer drug testing in animal models leading to failed clinical trials. A notable failed targeted therapy is saridegib (IPI-926), a Hedgehog pathway antagonist that increased survival in mouse models with malignant solid brain tumors^13^, but had no significant effect compared to placebo in patients with advanced chondrosarcoma participating in a Phase II randomized clinical trial^14^. Another immunomodulatory agent, TGN1412, was tested for safety in preclinical mice models and did not lead to toxicities in doses up to one hundred times higher than the therapeutic dose in humans^15^. However, when the drug advanced to Phase I testing, trial participants experienced multisystem organ failure and cytokine storm even with subclinical doses^16^. Anti-cancer vaccines have had similar issues in translating efficacy to human clinical trials. While therapeutic vaccines have successfully raised an immune response in mice, their effects in humans have been circumvented by immunological checkpoints and immunosuppressive cytokines that are absent in mice^17^. Examples of failed vaccines include Stimuvax, which had failed a non-small cell lung cancer phase III trial^18^, and Telovac, which failed in a pancreatic cancer phase III trial^19^.

The results of our study underscore the need for alternatives to classic cell culture and animal-based preclinical experiments. Human autopsy models have been used to test drugs in their early stages of development to mimic human physiological responses. *In silico* computer modeling may be a more accurate replacement to in vitro models, and involves implantation of cells onto silicon chips and using computer models to manipulate the cells’ physiologic response to agents and various parameters in the microenvironment^20^.

Given the track record of successful preclinical testing leading to failed clinical trials, efforts have been made to push forward direct testing in humans. In 2007, the European Medicines Agency and FDA proposed guidelines for bypassing preclinical testing and using micro-doses of therapeutic agents in humans^21^. The doses used in these “phase 0” studies are only a small fraction of the therapeutic dose, which are considered safe enough to bypass the usual testing required prior to phase I testing. Administering these micro-doses would help elucidate characteristics in drug distribution, pharmacokinetics, metabolism, and excretion in humans. Ideally, any new model that seeks to predict drug efficacy in cancer should be evaluated on the basis of its ability to predict clinical success and clinical failure. The widespread adoption of new preclinical models should ideally be accompanied by some measure of the model’s ability to predict clinical success as well as failure.

There were limitations to our study that should be acknowledged. Despite the large number of preclinical studies of lung cancer in the public domain, data on features of study design were inadequate. Analyses of cell culture data stratified on whether cells originated in lung cancer provided no indication that lung cancer cells constitute more predictive models; however, only 9 studies of approved drugs were conducted in cell lines of this type. Data on other features of cell and mouse models were too sparse to support even exploratory analysis of their predictive value. Another limitation is that some studies could not be included in the analysis owing to missing efficacy values. All of these were studies of failed drugs, and if efficacy values in the missing studies differed notably from those in studies included in our analysis, our results could obscure some true predictive value of the IC50 or TGI. However, notably different distributions of this nature seem unlikely, because all drugs – whether included or excluded for missing values – demonstrated a degree of preclinical efficacy that allowed them to advance to human studies. Regarding TGI efficacies, there were limitations in determining a standardized measure of efficacy for mouse models given the lack of standardized criteria on calculating drug effects in mice. The reported TGI values are based on raw tumor volumes extracted from tumor growth inhibition curves (if provided by articles) and applied to the equation as stated in the Methods, or reported TGI values derived from the same equation. A portion of articles used increase in life span as the measure of efficacy or a quantifiable effect on a molecular target, which were difficult to incorporate into the regression analysis used in this study and were thus excluded. Due to the naturally low proportion of approved compared to failed drugs, there is a sparse amount of data available for the former drug category, and thus any comparisons between the two drug classes may not be as robust. In addition, the approved drug category was lacking in immunotherapy agents as this study evaluated drugs in the pre-immunotherapy era.

It is important to note that when not accounting for the multiple studies per drug, we observed a significant association between efficacy values in Q3 and approval status, relative to values in Q1. There are three important points to note with these IC50 results. 1) In the cell experiments, we analyzed the data using two methods, one that accounts for the multiple studies per drug and one that ignores this characteristic of the data. Both methods have their limitations in this context and the truth likely lies between these two measures. 2) We would expect the relationship between drug approval and IC50 values to be characteristic of a monotonic relationship, meaning lower IC50 values correspond to greater odds of approval. In contrast to the individual quartile estimates, the trend statistics best capture the presence of this monotonic relationship, and in this study we should more heavily weigh the evidence from these statistics relative to the quartile measures. Both p trend statistics show the absence of a significant relationship between IC50 values and odds of drug approval. 3) Fig 1 and Fig 2 (A,B) agree with the absence of (or a weak) relationship between IC50 and approval status.

In conclusion, the findings of this study on pre-clinical testing of lung cancer therapies are consistent with prior concerns that cell and animal models are inadequate for identifying drugs that warrant human testing. Unfortunately, we found no evidence that either limiting in vitro models to cell lines derived from lung cancer or accepting narrower ranges of efficacy parameters is likely to improve performance of these conventional approaches. New models backed by evidence of their ability to predict clinical success and failure are needed.

## Acknowledgements

We thank Dr. Jorge Nieva and Dr. Victoria Cortessis for providing clinical and statistical guidance on this project.

## Supporting information

**Supplemental Table 1. List of Preclinical Cell Model References.** PubMed IDs (PMID) and Digital Object Identifiers (DOI) of articles that reported preclinical in vitro cell model experiments for all approved and failed lung cancer drugs, listed by drug name.

**Supplemental Table 2. List of Preclinical Murine Model References.** PubMed IDs (PMID) and Digital Object Identifiers (DOI) of articles that reported preclinical murine model experiments for all approved and failed lung cancer drugs, listed by drug name.

